# High dimensional single-cell analysis reveals iNKT cell developmental trajectories and effector fate decision

**DOI:** 10.1101/2020.05.12.070425

**Authors:** Thomas Baranek, Kevin Lebrigand, Carolina de Amat Herbozo, Loïc Gonzalez, Gemma Bogard, Céline Dietrich, Virginie Magnone, Chloé Boisseau, Youenn Jouan, François Trottein, Mustapha Si-Tahar, Maria Leite-de-Moraes, Thierry Mallevaey, Christophe Paget

## Abstract

CD1d-restricted invariant Natural Killer T (iNKT) cells represent a unique class of T lymphocytes endowed with potent regulatory and effector immune functions. Although these functions are acquired during thymic ontogeny, the sequence of events that give rise to discrete effector subsets remains unclear. Using an unbiased single-cell transcriptomic analysis combined with functional assays, we revealed an unappreciated diversity among thymic iNKT cells, especially among iNKT1 cells. Mathematical modelling and biological methods unravelled a developmental map whereby iNKT2 cells constitute a transient branching point towards the generation of iNKT1 and iNKT17 cells, which reconciles the two previously proposed models. In addition, we identified the transcription co-factor Four-and-a-half LIM domains protein 2 (FHL2) as a critical cell-intrinsic regulator of iNKT1 specification. Thus, these data illustrate the changing transcriptional network that guides iNKT cell effector fate.

## Introduction

Type I or invariant (i) Natural Killer T (iNKT) cells are a versatile population of thymus-derived αβT cells that play a critical role in the initiation and orchestration of immune responses in many pathological contexts including infection, cancer, inflammation and metabolic disorders (Bendelac et al., 2007; Godfrey et al., 2010). iNKT cells respond to lipid-based antigens presented by the quasi-monomorphic MHC class I-related CD1d molecule (Bendelac et al., 2007). Their swift response in the periphery is largely dependent on the existence of discrete pre-set subsets that secrete substantial amounts of either interleukin 4 (IL-4), IL-17A/F or interferon-γ (IFN-γ) akin to MHC-restricted T-helper cells and innate lymphoid cells (ILCs). This functional segregation is believed to be instigated in the thymus during their development involving many cues such as self-antigen recognition, transcription factors (TF), cytokine receptor signalling and cell-cell interactions.

The developmental steps underlying iNKT cell differentiation remain controversial. Two non-mutually exclusive models have been proposed. Initially, a linear maturation model has been reported (Benlagha et al., 2002; Pellicci et al., 2002) in which newly-selected stage 0 (CD24^+^) iNKT cells mature through stages 1 to 3 with the loss of CD24 expression and the sequential acquisition of CD44 and NK1.1. Functionally, iNKT cell maturation through these stages is associated with a reduced capacity to produce IL-4 and concomitant increase in IFN-γ production, which implies that IL-4-producing iNKT cells (comprised in stage 2) constitute an immature pool of cells. A more recent lineage differentiation model (Lee et al., 2013) suggests that discrete functional iNKT subsets develop from CD24^+^ iNKT0 (equivalent to stage 0), based on key TF expression into iNKT1 (IFN-γ^+^, T-bet^+^ PLZF^lo^), iNKT2 (IL-4^+^, PLZF^hi^) or iNKT17 (IL-17^+^, RORγt^+^ PLZF^int^). In this latter model, the three subsets appear to derive from a common CCR7^+^ intermediate progenitor (Wang and Hogquist, 2018). Although these findings constitute great strides, we still lack a clear understanding of the developmental steps governing iNKT cell development and their effector differentiation, at both the cellular and molecular levels.

Recent advances in genomic profiling have provided new insights into the highly divergent gene programs of thymic iNKT subsets (Engel et al., 2016; Georgiev et al., 2016; Lee et al., 2016). However, single cell approaches have used a limited number of cells and/or relied on iNKT0/1/2/17 subsets pre-sorted based on cell-surface markers. Although invaluable, these approaches precluded a comprehensive analysis of iNKT cell heterogeneity and possibly omitted functionally-relevant subsets or intermediate precursor cells.

In an attempt to further our understanding of the dynamics and checkpoints that dictate iNKT cell differentiation and sublineage commitment, we generated the transcriptomic profile of a large number of total thymic iNKT cells using single-cell (sc) RNA-sequencing in an unbiased fashion. In addition to the described iNKT0/1/2/17 subsets, unsupervised computational analysis of the transcriptomes uncovered previously unappreciated additional heterogeneity within iNKT cells, including several iNKT1 subsets. Moreover, by combining mathematical algorithms and biological assays, we propose a novel model for iNKT cell effector differentiation in which iNKT1 and iNKT17 subsets derive from iNKT2. Moreover, iNKT1 subsets arise linearly and sequentially from iNKT2 cells. Finally, we define a new molecular actor involved in iNKT1 effector fate. Thus, our data provide a comprehensive understanding of iNKT thymocyte heterogeneity and the transcriptional events that dictate sublineage decisions.

## Results

### scRNA-seq indicates substantial heterogeneity in iNKT thymocytes

To reveal the transcriptomic profile of iNKT thymocytes in an unbiased manner, we purified total iNKT cells (live CD3^+^ PBS57/CD1d tetramer^+^ cells) from juvenile (5 week-old) C57BL/6J mice (**Fig. S1)** and subjected them to droplet-based scRNA-seq. After control/filtering steps (**Fig. S1** and **Methods**), 3,290 cells were considered for further analyses. Unsupervised computational analysis, using the Louvain algorithm (Kiselev et al., 2017) indicated the existence of nine discrete clusters (I to IX) with different prevalence (**Fig. 1a**). Comparison of the expression pattern of genes in these communities (**Fig. 1b and Supplemental Table 1**) to published reports (Engel et al., 2016; Georgiev et al., 2016; Lee et al., 2016) indicated that most of these clusters matched with the previously identified iNKT0, iNKT1, iNKT2 and iNKT17 subsets. Differentially expressed genes (DEG) in cluster I (1.2 % of total iNKT cells) included *Lef1*, *Itm2a*, *Ccr9*, *Id3* and *Ldhb* (**Fig.1b-c and Supplemental Table 1**); several genes highly related to iNKT0 (Engel et al., 2016). Cluster II (2.5 % of total iNKT cells) was enriched for many genes involved in cytoskeletal control (*Stmn1*, *Ska1*, *Cfl1*), mitochondrial activity (*Atp5j*, *Atpif1*, *Cox5b*, *Cox6a1*), apoptosis regulation (*Birc5*, *Set*, *Sod1*) and mitosis (*Cenpe*, *Ccnb1*, *Cdk1*, *Cdc20*) (**Fig. 1b and Supplemental Table 1**) indicative of cell proliferation in either functional S or G2M phases (**Fig. 1d**) and was therefore named “cycling iNKT”. Clusters III (7.4 % of total) and IV (12.3 % of total) displayed high similarities characterized by a high expression of genes associated to iNKT2 biology such as *Zbtb16*, *Plac8* and *Icos* (**Fig.1b, 1e and Supplemental Table 1**) and were therefore referred to as iNKT2a and iNKT2b respectively (Engel et al., 2016; Lee et al., 2016). Cluster V (2.7 % of total) was homolog to iNKT17 cells as illustrated by the high expression of *Il1r1*, *Rorc*, *Ccr6, Tmem176a* and *Emb* (**Fig.1b, 1f and Supplemental Table 1**). iNKT1 cells were comprised within clusters VI-IX (73.7 % of total) based on their higher expression of *Id2*, *Nkg7*, *Klrb1c*, *Klrd1* and *Il2rb* (Engel et al., 2016) (**Fig. 1b, 1g and Supplemental Table 1**). As expected, the identified clusters showed differential and/or selective expression of some signature TFs such as *Rorc, Tbx21* and *Zbtb16* (Lee et al., 2013) (**Fig. 1h**). Altogether, this analysis indicates a previously unappreciated transcriptional heterogeneity in iNKT thymocytes.

**Figure 1:**
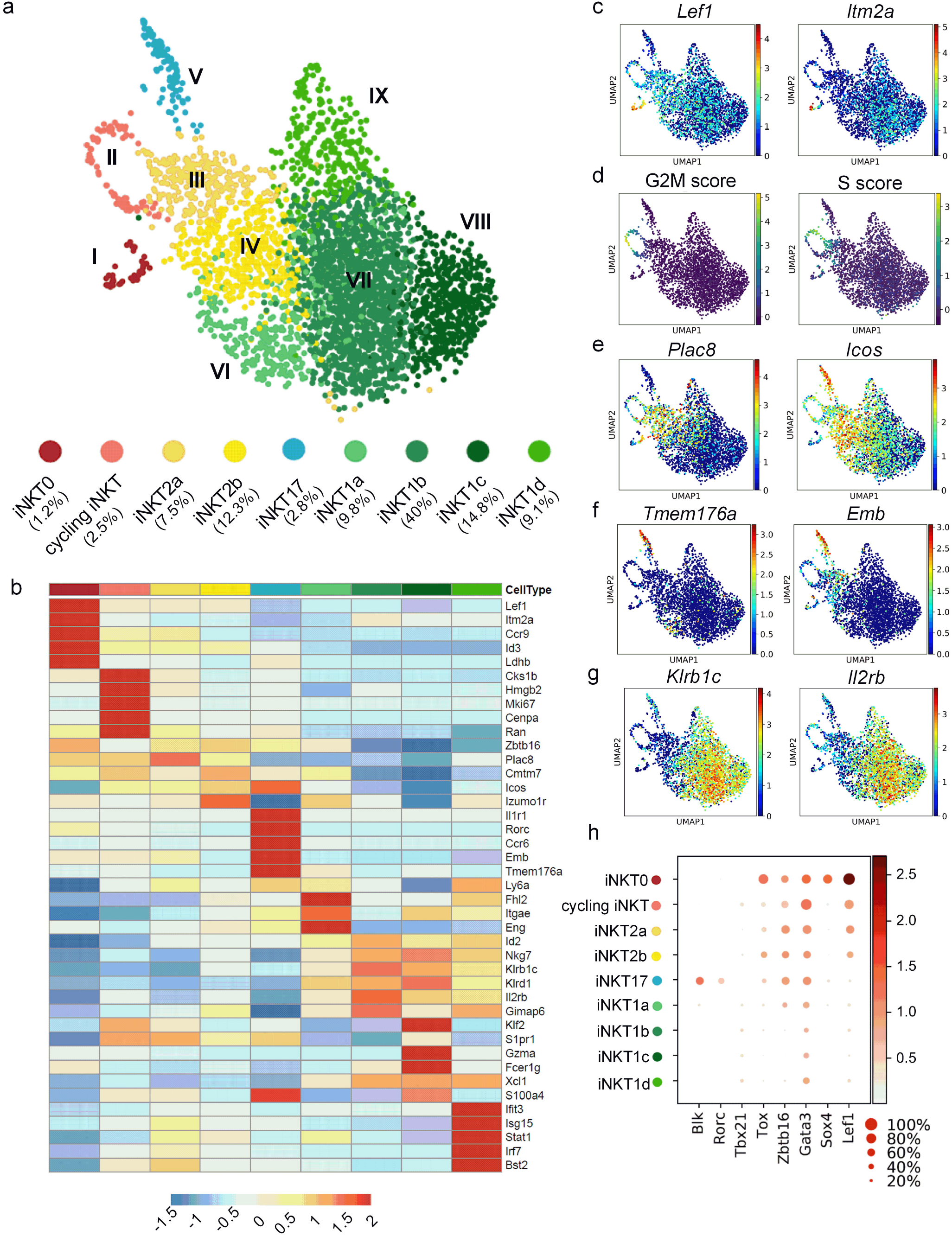
scRNA-seq identifies nine divergent thymic iNKT subsets. **a**, Identification of cell clusters in scRNA-seq data obtained from C57BL/6J total iNKT thymocytes using the graph-based Louvain algorithm (resolution = 1.5) on Uniform Manifold Approximation and Projections (UMAP). Each dot represents one cell (3,290 cells). **b,** Expression pattern of a representative list of cell subset-specific gene markers. **c, e-g**, Expression of selected gene markers in iNKT0 (**c**), iNKT2 (**e**), iNKT17 (**f**) and iNKT1 (**g**) identified in panel a. **d**, Representation of the quantitative scores for G2M and S phases for each cell. **h,** Dot map showing the expression of 8 selected genes encoding for key transcription factors in iNKT development per cluster. Color gradient and size of dots indicate gene expression intensity and the relative proportion of cells (within the cluster) expressing each gene respectively.

### Gene signature associated to iNKT thymocyte clusters

Newly-selected iNKT0 cells are believed to generate all effector subsets (Benlagha et al., 2005, 2002). Many genes encoding for proteins strongly associated with TCR signalling were enriched in iNKT0 cells (cluster I) such as *Pdcd1* (encoding for PD-1), *Cd27*, *Cd28*, *Slamf6* and *Cd81* (**Supplemental Table 1**). Several TF-encoding genes reported to control iNKT development were also upregulated in iNKT0 cells such as *Sox4* (Malhotra et al., 2018, p. 4), *Lef1* (Carr et al., 2015), *Id3* (Li et al., 2013) and *Tox* (Aliahmad and Kaye, 2008). Consistent with a role for CCR9 in thymocyte development and thymic trafficking (Uehara et al., 2002), *Ccr9* transcripts were up-regulated in iNKT0 cells (**Supplemental Table 1**). In addition, a large proportion of DEGs in iNKT0 encoded for products involved in nucleic acid regulatory processes and epigenetic landscape (**Supplemental Table 2**) suggesting that iNKT0 cells may undergo intense genome reprogramming.

In line with a role for miRNA in iNKT development (Henao-Mejia et al., 2013; Koay et al., 2016), *Drosha,* a key gene in miRNA biogenesis (Lee et al., 2003) was upregulated in both iNKT2 clusters (III and IV). Comparative analysis between iNKT2a and iNKT2b transcriptomes indicated that iNKT2a were enriched for genes related to tissue emigration such as *Klf2* and *S100a4* (Weinreich and Hogquist, 2008) (**Supplemental Table 3**) that may indicate the presence of potential iNKT thymic emigrants as recently proposed (Wang and Hogquist, 2018). *Ccr7* transcripts were also enriched in iNKT2a (Wang and Hogquist, 2018). Surprisingly, several DEGs typically associated with iNKT17 (Engel et al., 2016) were enriched in iNKT2a, including *Lmo4*, *Rora*, *Emb* and *Cxcr6* (**Supplemental Table 2**). This suggests that some cells within the iNKT2a cluster have iNKT17 potential. Similarly, iNKT2b cells (cluster IV) expressed higher levels of iNKT1 markers(Engel et al., 2016) than iNKT2a, such as *Klrb1c*, *Il2rb*, *Nkg7* (**Supplemental Table 3)**.

Among iNKT17 signature genes (**Supplemental Table 1**) were transcripts whose products are involved in tissue residency (*Itgb7*, *Ccr6*, *Ccr2*, *Aqp3*). In line with their anti-infectious role (Paget and Trottein, 2013), transcripts encoding for cytokine receptors were enriched in iNKT17 such as *Il17re*, *Il1r1*, *Il18r1* and *Il7r*. Interestingly, the percentage of mitochondrial RNAs (**Fig. S2a**) were enriched in iNKT17 suggesting particular energy requirements for the acquisition of iNKT17 effector functions. Analysis of mitochondrial mass and membrane potential confirmed the enhanced mitochondrial activity in iNKT17 thymocytes (**Fig. S2b**).

As previously mentioned, iNKT1 cells segregated into 4 clusters (VI to IX) with discrete gene programs (**Fig. 1b**). iNKT1a cells (cluster VI) presented many genes involved in RNA stability (*Npm1*, *Ubb*, *Cirbp*, *Pabpc1*, *Zfp36l2*, *Zfp36l1*) and translation activity (*Eif3e*, *Eif4a2*, *Eif3h*, *Eif1b*, *Eif1*, *Eef1b2*, *Eef1a1*, *Eef1g*, *Eef2*) (**Supplemental Table 1**). The iNKT1b cluster (cluster VII) comprised most of the iNKT1 cells (54.3%) and was characterized by the preferential expression of members from the GTPase of the immunity-associated protein (GIMAP) family (*Gimap1/3/4/5/6*) (**Supplemental Table 1**), a family of enzymes associated with TH1 differentiation and apoptosis regulation (Engel et al., 2016; Filén and Lahesmaa, 2010). Other genes associated with apoptosis regulation were also enriched in iNKT1b such as *Bcl2*, *Pdcd4* and *Cd44*. iNKT1b also displayed enriched expression of *Cxcr3* and *Itgb2* (encoding a subunit of LFA-1) that may indicate a pool of long-term resident cells. iNKT1c cells (cluster VIII) displayed a cytotoxic profile characterized by high expression of *Gzma/b*, *Ccl5* and many transcripts for Killer cell lectin type receptors such as *Klra1*, *Klra5*, *Klra7*, *Klra9*, *Klrc1*, *Klrc2*, *Klrd1*, *Klre1* and *Klrk1* (**Supplemental Table 1**). Finally, the iNKT1d subset (cluster IX) exhibited a unique signature featuring many interferon-stimulated genes such as *Ifit3*, *Isg15*, *Stat1* and *Irf7* (**Supplemental Table 1**). Hence, these data confirmed iNKT cell lineage and sublineage-specific transcriptomic profiles and unravelled new profiles reminiscent of differential functional activity.

### The clustering of cycling iNKT and iNKT1d is influenced by their biological state

Among the nine clusters, two of them displayed strong enrichment for genes associated to a particular biological state/function such as proliferation (cycling iNKT) and type I IFN response (iNKT1d) (**Supplemental Table 1**). Thus, we questioned whether their clustering was the consequence of particular biological states rather than a specific stage of differentiation. To address this possibility, sub-clustering of these two communities were analyzed in a semi-supervised manner using a gene set containing the top 50 DEGs in the other seven iNKT clusters as they may represent more iNKT-specific genes.

This resulted in the emergence of three new subclusters (0, 1 and 2) (**Supplemental Table 4a)** within cycling iNKT. Importantly, these subclusters were not associated with a particular phase of the cell cycle (**Fig. S3a)**. Rather, subclusters 0 and 1 presented high transcriptomic similarities with iNKT0, iNKT2b and iNKT2a, while subcluster 2 contained cells more related to iNKT1 clusters (**Fig. S3b**). In addition, monitoring of proliferative iNKT cells by means of mKi67 protein expression indicated that all effector subsets could cycle confirming the heterogeneous nature of cycling iNKT cells (**Fig. S3c**).

Subclustering of iNKT1d also generated three subclusters (**Supplemental Table 4b and Fig. S3d**) that resemble to either iNKT1 (cluster 0), iNKT2/17 (cluster 1) and iNKT2/1a/17 (cluster 2) (**Supplemental Fig. 3e**). This observation may suggest that type I IFN-signalling is involved in iNKT cell commitment/differentiation. Interestingly, *Ifnar1*^−/−^ mice displayed a reduction in iNKT1 and iNKT17 thymocyte numbers while iNKT0 and iNKT2 remained unaffected (**Fig. S3f**).

Altogether, these results suggest that once proliferation-related genes and ISG are removed, cycling iNKT and iNKT1d cells do not present discrete gene signatures and tend to merge with the other defined clusters. Since these two clusters appeared to be driven by their biological activity, they have been removed from the rest of our study.

### ScRNA-Seq reveals the developmental path of iNKT thymocytes

To assess similarities between clusters, we generated a clustermap using the Pearson’s method to order clusters by linkages based on similarity between the correlations (**Fig. 2a**). iNKT0 had the most unique signature albeit displaying some similarities with iNKT2a. As expected, iNKT1 and iNKT2 clusters formed two distinct groups (**Fig. 2a**). Of note, iNKT17 appeared closely related to iNKT2 (**Fig. 2a**). Within iNKT1 clusters, iNKT1a and iNKT1b displayed highest similarities (**Fig. 2a**).

**Figure 2:**
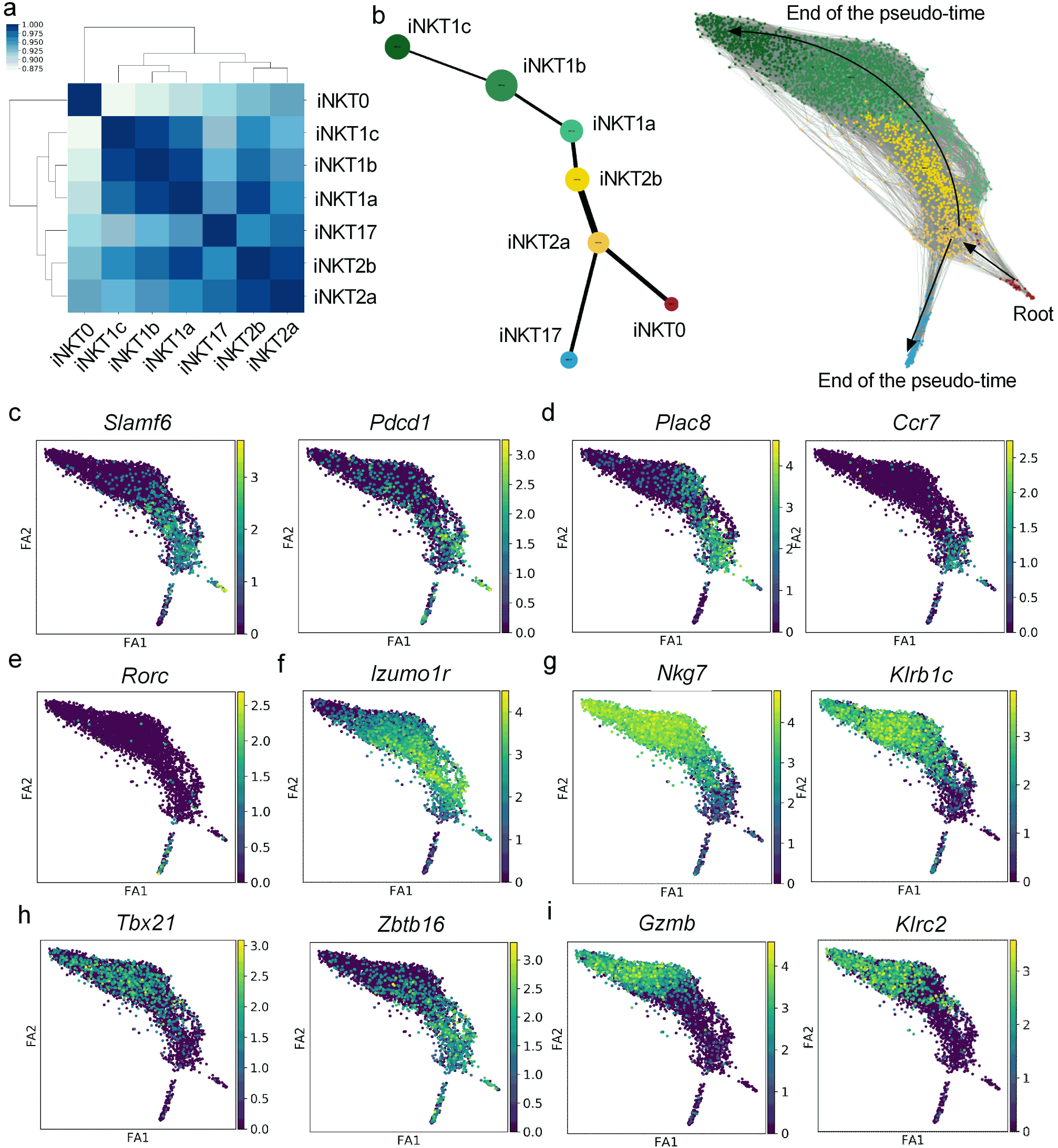
Predictive developmental trajectories of iNKT thymocytes. **a**, Clustermap of iNKT subsets comparing each pair of clusters using the Pearson’s correlation with hierarchical clustering. **b,** PAGA analysis applied to the scRNA-seq data from *Fhl2*^+/+^ iNKT thymocytes. PAGA graph (left panel) and PAGA-initialized single-cell embedding (right panel) are shown. Weighted edges represent a statistical measurement of connectivity between clusters. **c-i**, Visualization of selected markers on PAGA-initialized single-cell embedding alongside the trajectories defined in panel b.

To investigate the dynamics and trajectories underlying the iNKT cell differentiation program, we attempted to create a pseudo-time ordering of iNKT cell transcriptomes to map progenitors to fates. The use of partition-based graph abstraction (PAGA) (Wolf et al., 2019), in an unsupervised manner, enabled to generate a graph-like-map based on connectivity between clusters and single-cell embedding (**Fig. 2b**). Based on the literature, iNKT0 cells were defined as the root of the pseudo-time ordering. This could be illustrated by the expression of early markers of iNKT positive selection such as *Slamf6* and *Pdcd1* (Lu et al., 2019) (**Fig. 2c).** In the predicted model, the PAGA path suggested that iNKT0 cells gave rise to iNKT2a cells (**Fig. 2b)**. This transition is accompanied by a strong induction of *Plac8* and *Ccr7* (**Fig. 2d**), in line with their migration from the cortex to the medulla (Wang and Hogquist, 2018). Then, iNKT2a appeared as a node that split into two branches leading to either iNKT17, as illustrated by *Rorc* expression (**Fig. 2b and 2e**) or iNKT2b, which was associated with the upregulation of *Izumo1r* (**Fig. 2b and 2f**). Importantly, emergence of several iNKT1 marker genes such as *Nkg7* and *Klrb1c* (Engel et al., 2016) (**Fig. 2g**) were found in iNKT2b. Strikingly, iNKT2b appeared to differentiate towards iNKT1 cells with the iNKT1a cluster as the earliest defined iNKT1 subset (**Fig. 2b**). Pseudo-time progression within the iNKT1 branch led to the sequential emergence of iNKT1b and iNKT1c clusters (**Fig. 2b**), illustrated by *Tbx21* upregulation and paralleled with a loss of *Zbtb16* (**Fig. 2h**), an important step for iNKT1 differentiation (Pobezinsky et al., 2015). iNKT1c emerged as a terminal stage of differentiation within the iNKT1 sublineage associated with high expression of *Gzmb* and *Klrc2* (encoding for NKG2C) (**Fig. 2i**). Thus, this model predicts a cascade of molecular events that parallel and potentially drive the differentiation of effector iNKT cells in which iNKT2 clusters appear poised at the crossroad of iNKT sublineages.

### Identification of thymic iNKT subsets based on cell-surface markers

Building on the scRNA-Seq data, we defined a set of surface molecules to track and biologically evaluate these newly-defined thymic iNKT subsets. Based on CD24 (*Cd24a*), Sca-1 (*Ly6a*), FR4 (*Izumo1r*), CD138 (*Sdc1*) and NK1.1 (*Klrb1c*) (**Fig. 3a**), we were able to identify iNKT0 (CD24^+^, Sca-1^−^, NK1.1^−^), iNKT2a (CD24^−^, NK1.1^−^, CD138^−^, FR4^−^), iNKT2b (CD24^−^, NK1.1^−^, CD138^−^, FR4^+^), iNKT17 (CD24^−^, Sca-1^hi^, CD138^+^), iNKT1a (CD24^−^, Sca-1^hi^, CD138^−^, NK1.1^dim^), iNKT1b (CD24^−^, Sca-1^+^, CD138^−^, NK1.1^hi^) and iNKT1c (CD24^−^, Sca-1^−^, CD138^−^, NK1.1^hi^) (**Fig. 3b**). We compared our gating strategy to the classical maturation stages based on expression of CD44 and NK1.1. As expected, iNKT0 were CD24^+^, NK1.1^−^ and CD44^-/low^ (stage 0/1) (**Fig. 3c**). Stage 2 (CD24^−^CD44^+^NK1.1^−^) cells comprised both iNKT2 subsets as well as iNKT17 cells, some of which can express low levels of NK1.1 (**Fig. 3c**). Notably, iNKT2b displayed a slightly brighter expression of CD44 compared to iNKT2a (**Fig. 3c**). All subsets of iNKT1 displayed a stage 3 phenotype (CD44^+^ NK1.1^dim/+^) (**Fig. 3c**).

**Figure 3:**
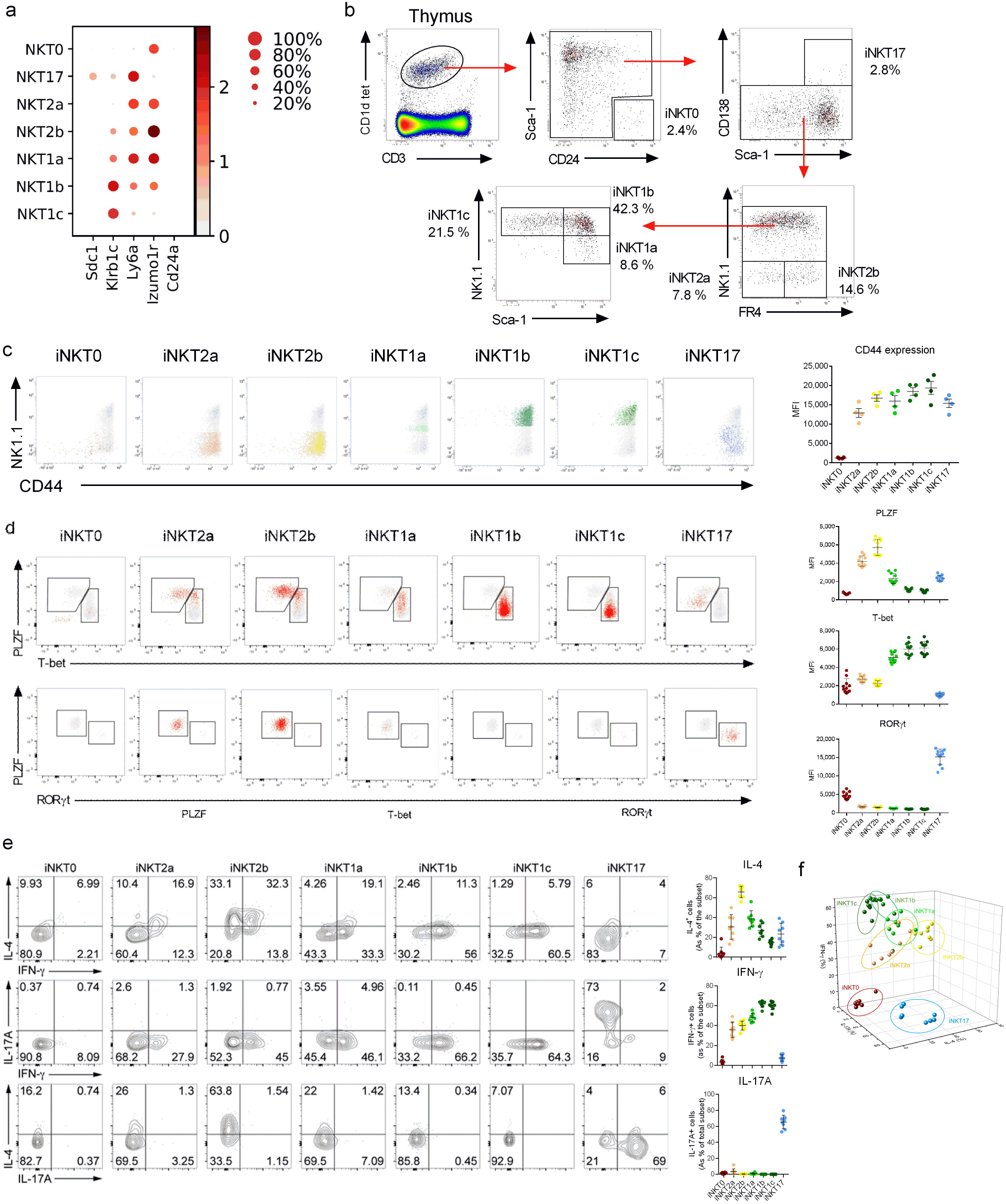
Identification of iNKT clusters using cell surface markers. **a,** Dot map showing the expression of 5 surface markers per cluster versus all others. Color gradient and size of dots indicate gene expression intensity and the relative proportion of cells (within the cluster) expressing each gene respectively. **b**, Gating strategy for the identification of the iNKT subsets in the thymus of 5 week-old C57BL/6J mice. **c,** Representative overlay dot plots showing CD44 and NK1.1 expression in iNKT subsets as obtained in (b) is shown in the left panel. Individual values and means ± SEM of mean fluorescence intensity (MFI) of CD44 for each subsets are depicted in the right panel. Data are representative of one experiment out of three. **d,** Flow cytometry showing PLZF, T-bet (upper panel) and RORγt within PLZF^+^ T-bet^−^ cells (middle panel) expression in iNKT subsets as identified in panel b. Data are representative of three independent experiments. Individual values and means ± SEM of mean fluorescence intensity (MFI) for each subsets are depicted in the right panel. **e**, IFN-γ, IL-17A and IL-4 production in PMA/ionomycin-stimulated iNKT subsets as identified in panel b. Of note, iNKT17 were defined as RORγt^+^ due to down-modulation of Sdc-1 upon stimulation. Data are pooled from two independent experiments. Individual values and means ± SEM for each subset are depicted in the right panel. **f**, 3D scatter plot for cytokine production (IL-4, IFN-γ and IL-17A). Each data point represent one mouse (n = 8). Data are pooled from two independent experiments.

The differential expression of PLZF, T-bet and RORγt (Lee et al., 2013) also confirmed our assignment. iNKT0 cells were PLZF^−^, T-bet^−^ and RORγt^dim^ (**Fig. 3d**). Both iNKT2 subsets expressed high levels of PLZF although its expression was slightly elevated in iNKT2b (**Fig. 3d**). Of note, iNKT2 subsets expressed low levels of T-bet and no RORγt (**Fig. 3d**). iNKT17 cells expressed low levels of T-bet, intermediate levels of PLZF and the highest levels of RORγt (**Fig. 3d**). All iNKT1 subsets expressed T-bet albeit at various intensities (**Fig. 3d**). In line with our proposed model of differentiation, a gradual decrease in PLZF expression was paralleled with T-bet acquisition in iNKT1 subsets (**Fig. 3d**).

We also tested the cytokine profile of these subsets of iNKT thymocytes upon short-term *in vitro* stimulation with PMA/ionomycin. As expected, both iNKT2 subsets produced IL-4 although iNKT2b produced higher levels as compared to iNKT2a (**Fig. 3e**). iNKT17 almost exclusively produced IL-17A and iNKT1 subsets produced IFN-γ (**Fig. 3e**). However, in line with a previous report (Cameron and Godfrey, 2018), we observed that *bona fide* iNKT1 - especially iNKT1a - and iNKT17 cells could produce IL-4 at some extent (**Fig. 3e**). Consistent with the proposed developmental trajectories, a decrease in IL-4 production capacity among iNKT1 cells paralleled their maturation/differentiation (iNKT1a > iNKT1b > iNKT1c) (**Fig. 3e**). In line with previous reports (Benlagha et al., 2002; Pellicci et al., 2002), both iNKT2a and iNKT2b subsets co-produced IFN-γ and IL-4, but no detectable IL-17A (**Fig. 3e**). Moreover, three-dimensional plotting indicated that iNKT subsets clustered based on cytokine profile (**Fig. 3f**). This analysis suggested that the iNKT2b and iNKT1a are relatively close in terms of cytokine profile (**Fig. 3f**) and a progression towards a Th1-like profile is observed in iNKT1 subsets **(Fig. 3f**). Thus, the 7 clusters defined by scRNA-Seq can be readily identified by flow cytometry using cell surface markers and have discrete cytokine signature.

### Multiple iNKT subsets have the potential to egress the thymus

The nature of thymus-emigrating iNKT cells remains controversial and both iNKT2 (Wang and Hogquist, 2018) and iNKT17 (Milpied et al., 2011) have been shown to leave the thymus. Based on our transcriptomic data, we compared the preferential expression of egress/retention gene markers between clusters (**Fig. 4a**). iNKT2a, iNKT17 and iNKT1c clusters were enriched for *Klf2*, a master transcriptional regulator of sphingosine 1-phosphate receptor (Weinreich and Hogquist, 2008) (**Fig. 4a and Supplemental Table 1**) that may indicate at least several functional subsets of iNKT thymic emigrants. In addition, several genes of the KLF2 regulon such as *Emp3*, *S100a4* and *S100a6* displayed a similar pattern of expression (**Fig. 4a**). Of note, the expression of *Ccl5*, a regulator of *Klf2* expression (Kroetz and Deepe, 2011) was also found elevated in iNKT1c (**Supplemental Table 1**). Conversely, iNKT0 expressed CD69 while iNKT2b and iNKT1a/b subsets expressed *Cxcr3* (**Fig. 4a**), two genes encoding for proteins involved in thymic retention (Drennan et al., 2009; Weinreich and Hogquist, 2008). To test these observations *in vivo*, we attempted to block thymocyte egress by means of FTY720, a selective sphingosine 1-phosphate receptor agonist that induces its internalization (Mandala et al., 2002). Prolonged FTY720 treatment led to a significant increase in iNKT thymocytes (**Fig. 4b**). This artificial retention led to a specific enrichment for iNKT2, iNKT17 and iNKT1c subsets (**Fig. 4c**). Thus, these data may suggest that these three sublineages of functional iNKT cells have the potential to leave the thymus.

**Figure 4:**
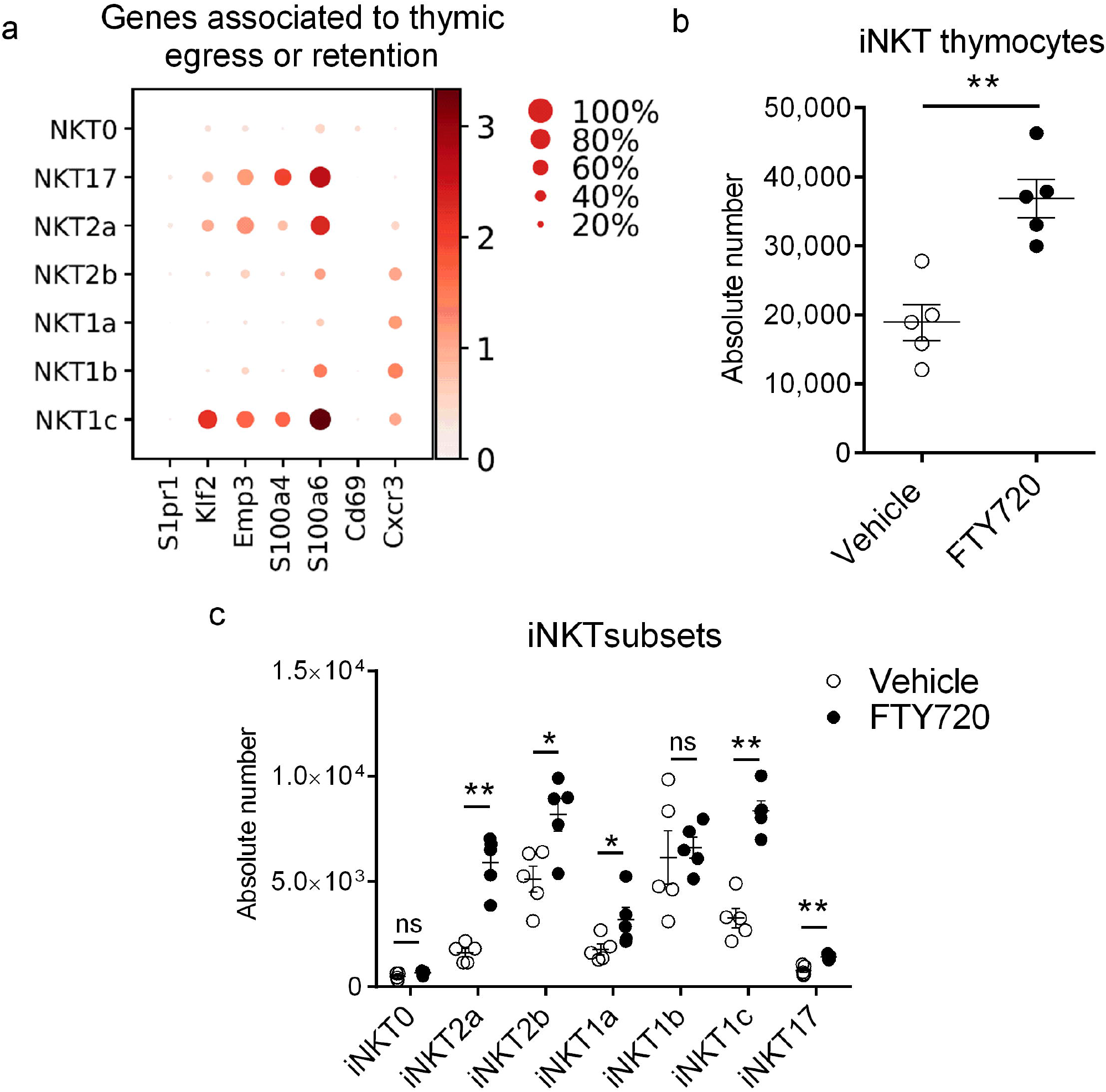
Transcriptomic and biological evaluation of iNKT thymocyte egress. **a**, Dot map showing selected gene expression per cluster versus all other cells. Color gradient and size of dots indicate gene expression intensity and the relative proportion of cells (within the cluster) expressing each gene respectively. **b-c,** Mice received daily vehicle (○) or FTY720 (•) by the i.p. route during 7 days. Mice were sacrificed on day 8 and thymi were harvested. The absolute number of iNKT cells **(b)** and subsets **(c)** were determined by flow cytometry. Individual values and means ± SEM of one representative experiment out of two is shown (n = 5 mice/group).*, p<0.05; **, p<0.01.

### iNKT2 subsets can generate both iNKT1 and iNKT17 cells

To biologically evaluate our predictive developmental model, various purified iNKT subsets were cultured with *Traj18*^−/−^ total thymocytes in presence of IL-7 and IL-15, two critical factors for iNKT cell survival and differentiation (Matsuda et al., 2002). After 4 days of culture, iNKT2a cells generated iNKT2b, iNKT17 and iNKT1 (**Fig. 5a**) but no iNKT0 cells (not shown). Of note, only 12 % of iNKT2a cells retained their original profile suggesting these cells may be in a transient stage of differentiation (**Fig. 5a**). In line with the predicted computational model (**Fig. 2a**), iNKT2b generated iNKT1 but few iNKT17 as compared to iNKT2a (**Fig. 5a**) suggesting that this very subset mainly lost its plasticity at least in this *ex vivo* setting. As expected, iNKT1a and iNKT1b generated large amounts of iNKT1 cells (**Fig. 5a**). However, iNKT1a could still generated a substantial amount of iNKT2 (**Fig. 5a**). Importantly, this effect was much more limited with iNKT1b cells (**Fig. 5a**). Consistent with their proliferative and pluripotent nature (**Fig. 2b and Fig. S3c**), culture with iNKT2a resulted in the generation of higher number of iNKT cells compared to others subsets (**Fig. 5b**).

**Figure 5:**
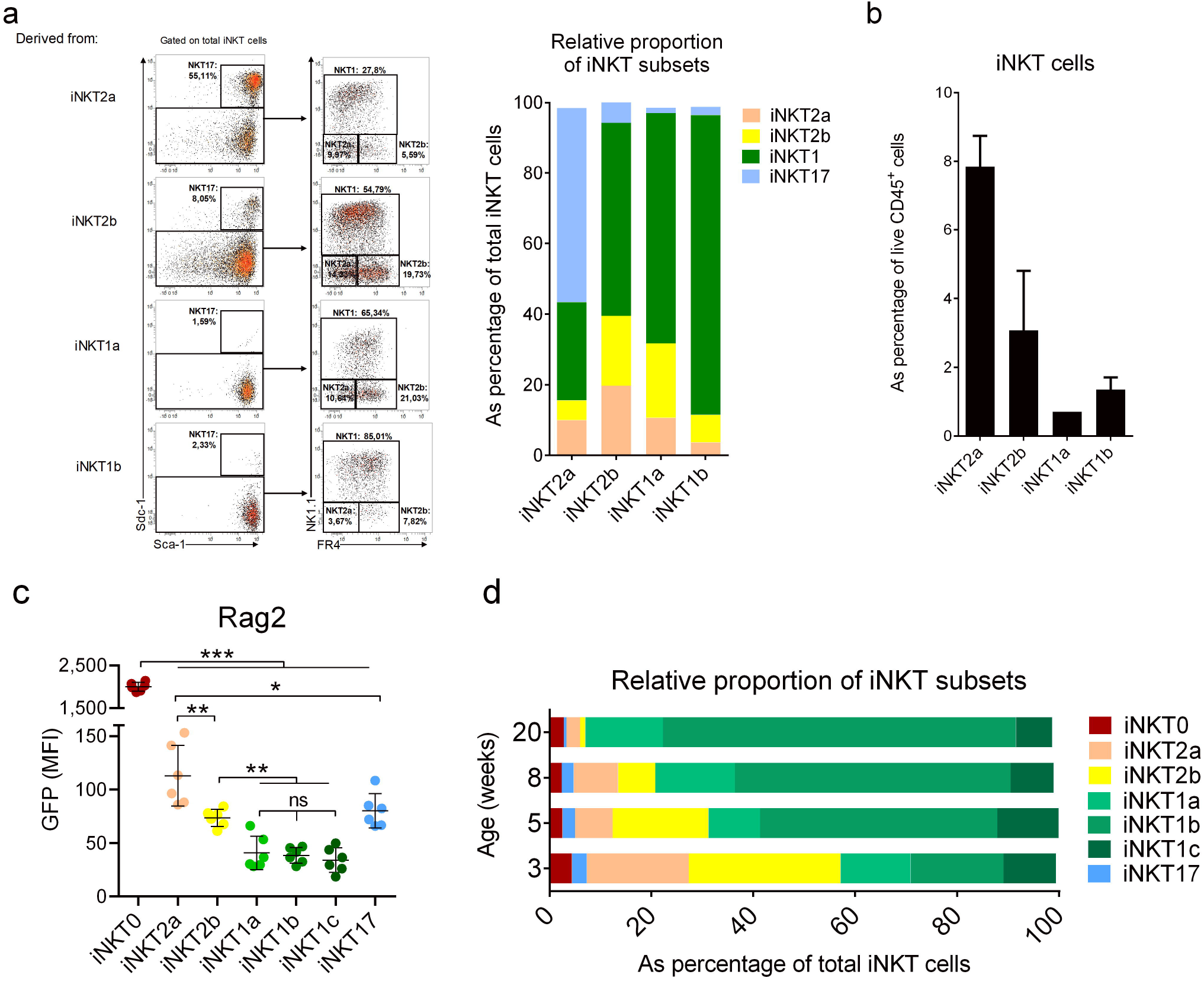
iNKT1 and iNKT17 effector cells derive from iNKT2 precursors. **a-b,** Purified iNKT subsets were cultured for 4 days with total thymocytes from *Traj18*^−/−^ mice in presence of IL-7 and IL-15. **a,** Representative dot plots of generated iNKT cells from indicated subsets are depicted in the left panel. Mean of iNKT subsets’ relative proportion obtained from three independent experiments is shown in the right panel. **b,** Percentage of total iNKT cells generated within live cells after 4 days of culture was determined by flow cytometry. Mean ± SEM of three independent experiments is shown. **c,** Flow cytometry of GFP in iNKT subsets using *Rag2*^GFP^ mice. Representative data of two independent experiments are shown in the upper panel. Individual values and means ± SEM of MFI of GFP are shown. ns, not significant; *, p<0.05; **, p<0.01, ***, p<0.001. **d,** Relative proportion of iNKT subsets according to the indicated age from littermate mice. Mean pooled from two independent experiments is shown (n = 8).

To monitor time to positive selection of iNKT cells, we took advantage of Rag2^GFP^ mice by following the GFP decay as an indicative marker of time from positive selection (Boursalian et al., 2004). iNKT0 cells expressed the highest level of GFP indicating recent positive selection (**Fig. 5c**). In line with the maturation process, levels of GFP varied between subsets with intermediate levels in iNKT2 and iNKT17 cells but low levels in iNKT1 subsets (**Fig. 5c**). A gradual reduction was observed between iNKT1 subsets reinforcing the possibility that iNKT1a cells were an immature stage in iNKT1 differentiation (**Fig. 5c**).

Since at least some iNKT2 thymocytes appeared as intermediate of development in iNKT effector fate, we assessed how iNKT subset prevalence evolve with age. While iNKT2 cells were the major subsets in 3-week-old thymi, we observed a striking decrease in their frequency as mice aged (Cruz Tleugabulova et al., 2019; Lee et al., 2013) (**Fig. 5d**). Consistent with their putative thymic resident nature, this decrease was paralleled with a strong enrichment for iNKT1b (**Fig. 5d**). No major modulation was observed for iNKT0, iNKT1a and iNKT1c (**Fig. 5d**). Of note, relative proportion of iNKT17 cells was severely decreased in 20-week-old mice (**Fig. 5d**). Altogether, these biological approaches confirm the predictive RNA-based model and indicates that most iNKT2 are pivotal in iNKT commitment towards iNKT1 and iNKT17 profiles.

### *Fhl2* controls commitment to iNKT1 lineage

iNKT1a cells appeared as the earliest committed iNKT1 cells (**Fig. 2a**) and therefore it is possible that some of the signature genes within this cluster may be involved in effector fate specification. Among those candidates is *Fhl2* (**Fig. 1b and Supplemental Table 1**), a gene encoding for the transcription co-factor Four and a half LIM domain 2 (FHL2) recently shown to control NK cell development (Baranek et al., 2017) but whose function in T cell biology remains elusive. We found that *Fhl2*^−/−^ mice had a decrease in total thymic iNKT cell numbers (**Fig. 6a**). Although a slight decrease in total thymocyte number was noted (**Fig. S4a**), no clear modifications were observed neither on DN subsets nor on CD4SP, CD8SP and DP (**Fig. S4b**). Based on TF expression, we observed that *Fhl2*^−/−^ iNKT thymocytes displayed a reduced number of iNKT1 compared to their *Fhl2*^+/+^ counterparts while both iNKT2 and iNKT17 remained unaffected (**Fig. 6b**). Selective iNKT1 cell defects were also observed in peripheral tissues of FHL2-deficient mice (**Fig. S5**). Using our staining strategy, we observed that the lack of iNKT thymocytes in *Fhl2*^−/−^ mice was restricted to iNKT1b and iNKT1c (**Fig. 6c**). Conversely, the number of iNKT2b cells was slightly increased in *Fhl2*^−/−^ mice (**Fig. 6c**) suggesting a block at the iNKT2b to iNKT1a transition phase.

**Figure 6:**
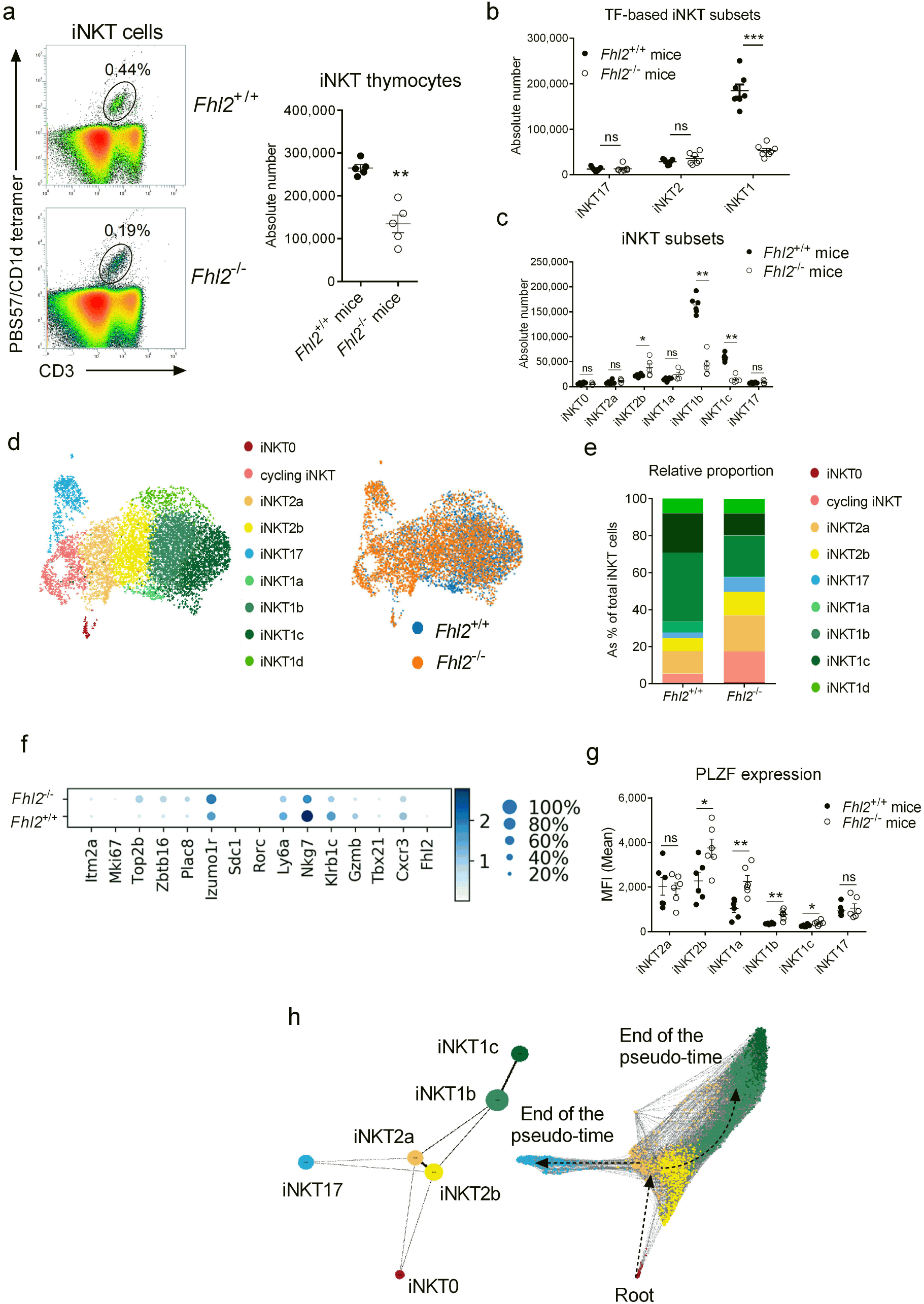
*Fhl2* deficiency affects iNKT cell development. **a,** Flow cytometric analyses of iNKT thymocytes in *Fhl2*^+/+^ and *Fhl2*^−/−^ littermate mice. Representative dot plots from eight experiments are shown in the left panel. Individuals and means ± SEM of absolute number of iNKT thymocytes from one representative experiment out of eight are shown in the right panel. **, p<0.01. **b,** Flow cytometry of iNKT subsets from *Fhl2*^+/+^ or *Fhl2*^−/−^ mice based on PLZF and T-bet expressions: iNKT17 (PLZF^dim/-^ T-bet^−^), iNKT2 (PLZF^hi^ T-bet^−^) and iNKT1 (PLZF^dim/-^ T-bet^+^). Data are representative of two independent experiments. Individual values and means ± SEM for each subsets are depicted. **c**, Flow cytometry analysis of iNKT subsets in thymi of *Fhl2*^+/+^ and *Fhl2*^−/−^ mice. Individual values and means ± SEM for each subsets from two independent experiments are shown. ns, not significant; ***, p<0.001. **d,** Aggregated UMAP representation of scRNA-seq data from *Fhl2*^+/+^ (3,290 cells) and *Fhl2*^−/−^ (4,189 cells) iNKT thymocytes. Clusters identities are shown in the left panel and WT onto KO projection is shown in the right panel. **e,** Relative proportion of iNKT clusters in *Fhl2*^−/−^ and *Fhl2*^+/+^ mice from scRNA-seq data. **f,** DEG in *Fhl2*^+/+^ vs *Fhl2*^−/−^ iNKT thymocytes from scRNA-seq analysis. Dot map depicting selected gene expression per cluster is shown in the lower panel. Color gradient and size of dots indicate gene expression intensity and the relative proportion of cells (within the cluster) expressing each gene respectively. **g**, Flow cytometry of PLZF expression in *Fhl2*^+/+^ and *Fhl2*^−/−^ iNKT subsets. Individual values and means ± SEM of PLZF MFI for each subsets from one experiment out of three are depicted. ns, not significant; *, p<0.05; **, p<0.01. **h**, PAGA analysis applied to the scRNA-seq data from *Fhl2*^−/−^ iNKT thymocytes. PAGA graph (left panel) and PAGA-initialized single-cell embedding (right panel) are shown. Weighted edges represent a statistical measurement of connectivity between clusters.

To evaluate this at the transcriptomic scale, we generated scRNA-seq data from *Fhl2*^−/−^ iNKT thymocytes. By applying graph-based clustering of *Fhl2*^−/−^ iNKT after aggregation of WT transcriptomes, we confirmed the presence of most of the 9 original clusters found in WT mice, with the exception of iNKT1a, which were absent from *Fhl2*^−/−^ iNKT cells, suggesting the critical role of this gene in the transcriptomic regulation of this cluster (**Fig. 6d,e**). Consistent with the biological data, a reduced proportion within iNKT1 clusters was observed in *Fhl2*^−/−^ iNKT transcriptomes (**Fig. 6e**). Paralleling their higher relative proportion, several key iNKT2 marker genes were enriched in *Fhl2*^−/−^ iNKT cells including *Zbtb16, Izomu1r* and *Plac8* (**Fig. 6f and Supplemental Table 5a**). However, cluster-by-cluster analysis did not show any significant modulation for *Zbtb16* (**Supplemental Table 5b-f**). Post-transcriptional regulation of PLZF is required to allow iNKT1 specification(Pobezinsky et al., 2015). In line, the levels of PLZF protein were significantly increased in *Fhl2*^−/−^ iNKT subsets at the exception of iNKT2a and iNKT17 (**Fig. 6g**). Compared to controls (**Fig. 2b**), application of the PAGA algorithm on *Fhl2*^−/−^ iNKT transcriptomes revealed some disruptions in the interconnectivity between clusters including the iNKT2/iNKT1 transition (**Fig. 6h**). Collectively, FHL2 controls iNKT1 lineage commitment possibly through its post-transcriptional regulation of PLZF.

## Discussion

Using high dimensional scRNA-seq and biological assays, we herein characterize the cascade of events occurring in the aftermath of iNKT thymocyte positive selection. This analysis enabled to highlight an unexpected heterogeneity, many of those subsets being transitional stages linked to the acquisition of their effector program. Importantly, we were able to analyze these transient stages through pseudo-time ordering and *ex vivo* differentiation assays.

This reveals a continuous and dynamic process experienced by differentiating iNKT thymocytes following positive selection. In this process, iNKT0 matured to uncommitted iNKT2a cells characterized by expression of *Plac8* and *Ccr7*. At the protein level, this transition phase is accompanied by a strong expression of PLZF. Interestingly, the transcriptional program of iNKT2a is reminiscent to a cluster of central and transitional Mucosal-Associated Invariant T (MAIT) thymocytes (Koay et al., 2019; Legoux et al., 2019). Thus, iNKT2a likely represent a precursor stage preceding effector fate commitment as previously suggested (Wang and Hogquist, 2018). In agreement with this, iNKT2a emerged as a pivotal subset that can either egress from the thymus or fulfil their differentiation program. As such, iNKT2a can commit to the iNKT17 or iNKT1 sublineages. Interestingly, commitment to the iNKT1 effector fate seems to require transitional states that closely resemble to iNKT2 (*e.g*. iNKT2b and iNKT1a). This raises the question whether iNKT2 have to be considered as an intermediate, a terminal stage of differentiation or a combination of both. In our *ex vivo* differentiation model, consistent with pioneer studies (Benlagha et al., 2002; Pellicci et al., 2002) neither iNKT2a nor iNKT2b cells maintained their original stage, although we cannot exclude that a small proportion of these latter can retain a long-term iNKT2 profile. Moreover, we constantly observed that most of the iNKT2 thymocytes co-produced IL-4 and IFN-γ. It is noteworthy that murine MAIT thymocytes do not appear to differentiate into *bona fide* MAIT2 cells (Legoux et al., 2019). In any case, iNKT2 appear to conserve a certain degree of plasticity that enables them to generate either iNKT1 or iNKT17 cells, a situation which is less obvious for these two latter subsets. In addition, iNKT2 numbers are strongly reduced with age. Although this favours the hypothesis of a developmental precursor, it can also result from changes in the thymic environment (*e.g.* cytokine and ligand availability) that may interfere in the iNKT2 differentiation process. Further works will be required to delineate the true nature of iNKT2 in iNKT cell effector fate. Our data also pointed towards an unappreciated transcriptomic heterogeneity within the pool of iNKT1 cells. First, iNKT1a emerges as a pivotal subset in iNKT1 specification in which the transcription co-factor FHL2 is involved. Its recognized repressive activity on PLZF (Tran et al., 2016) may, at least in part, explains the role of FHL2 in iNKT1 effector fate (Pobezinsky et al., 2015). Similarly, the mechanisms that regulate FHL2 expression and activity in iNKT cells deserve further consideration. Moreover, it will be interesting to test the cell-intrinsic involvement of FHL2 in other PLZF-dependent immune cells such as ILC, MAIT or γδT subsets. Importantly, FHL2 has been shown to interact with several key factors in lymphocyte development such as Id2, Id3 and Nur77 (Tran et al., 2016). Thus, the possibility that FHL2 regulates various innate and/or adaptive lymphocytes will be worth investigating.

It is noteworthy that FHL2 deficiency does not fully abrogate iNKT1 differentiation suggesting that commitment towards iNKT1 lineage may be under several and potentially redundant layers of regulation. Accordingly, a study elegantly demonstrated that PLZF silencing during iNKT1 commitment was under the control of a family of miRNAs (Let-7) (Pobezinsky et al., 2015). Our data also indicate that multiple subsets of iNKT cells can naturally egress the thymus. These encompass both multipotent iNKT cells (iNKT2a) and fully-differentiated subsets (iNKT1 and iNKT17). This may provide a strong advantage for the host in response to a peripheral insult such as infection by having on-site both iNKT subsets that can readily respond as well as iNKT precursors that can differentiate on-demand according to the inflammatory context.

Other functional iNKT subsets have been identified in the periphery such as iNKT10 (Sag et al., 2014) and iNKTFH (Chang et al., 2011). However, we did not evidence such subsets based on our sc-RNA-seq analysis that mainly supports extrathymic requirements. In agreement, a recent study demonstrated that long-term activation was required to observe the emergence of such functions irrespective of the subset (Cameron and Godfrey, 2018).

Altogether, we propose a novel model for the development of effector iNKT cells. Our single cell approach highlights the dynamic sequence of transcriptional events that illustrates how iNKT cell can reach such a wide variety of specificities. Thus, some of the genes described here and modulated during this developmental pathway will likely constitute candidate molecules in iNKT cell biology with potential impact in the context of infection, cancer and inflammation.

## Abbreviations

DEG: differentially-expressed gene
FHL2: Four-and-a-half LIM domain 2
iNKT: invariant Natural Killer T cell
IL: interleukin
IFN: interferon
PAGA: partition-based graph abstraction
TCR: T cell receptor
TF: transcription factor
WT: wild-type

## Acknowledgements

This work was supported by the Agence Nationale de la Recherche “JCJC program” (ANR-19-CE15-0032-01) and by the Region Centre-Val de Loire under the program “ARD 2020 Biomédicaments” (Project 7UP). M.S., and C.P. are supported by Inserm. F.T. and M.L-d-M. are supported by CNRS. T.B., L.G., and C.B. are supported by the University of Tours. Roxane Lemoine (BCR Facility, University of Tours) and Julie Cazareth (CNRS, IPMC, Sophia-Antipolis) are acknowledged for their assistance with cell sorting. We acknowledge the UCAGenomiX platform, partner of the National Infrastructure France Génomique, supported by the Commissariat Aux Grands Investissements (ANR-10-INBS-09-03, ANR-10-INBS-09-02). M. Kronenberg (La Jolla Institute for Allergy & Immunology) and P. Fink (University of Washington) are acknowledged for the kind gift of *Traj18*^−/−^ and *Rag2*^GFP^ mice respectively. We also thank the University of Tours animal facility for excellent mouse husbandry and care. We thank the NIH tetramer core facility (Emory University) for providing CD1d tetramers.

## Author contributions

KL designed and performed the bioinformatic analysis and analyzed data. C.d.A.H., G.B., C.D., L.G., C.B., F.C and Y.J. performed the experiments. F.T., P.B., M.S., M.L-d-M. and T.M. provided key materials and critical input. T.B. and C.P. supervised the project, performed experiments, analyzed data and wrote the paper with the input of all authors.

## Declaration of interests

The authors declare no competing interests

Further information and requests for resources and reagents should be directed to and will be fulfilled by the Lead Contact, Christophe Paget.

## Methods

### Mice

Wild-type (WT) C57BL/6JRj (B6) mice were purchased from Janvier Labs (Le Genest-St-Isle, France). B6 *Fhl2*^−/−^, *Ifnar1*^−/−^, *Traj18*^−/−^ and *Rag2*^GFP^ mice mice have been previously described(Boursalian et al., 2004; Chu et al., 2000; Moran et al., 2011; Müller et al., 1994),(Chandra et al., 2015). Experiments using *Fhl2*^−/−^ mice were confirmed using *Fhl2*^−/−^ and *Fhl2*^+/+^ littermates obtained from intercrossing of *Fhl2*^+/−^ mice. Mice were bred and maintained under specific pathogen-free conditions in-house and were used at 5-8 weeks of age. All animal experimentation was performed according to the national governmental guidelines and was approved by our local and national ethics committee (CEEA.19, #201604071220401-4885).

### Reagents and antibodies

Cell surface staining was performed using antibodies from BD Biosciences (San Diego, CA, USA), Biolegend (San Diego, CA, USA) and eBioscience (San Diego, CA, USA): APC-eFluor780-conjugated anti-CD45 (30-F11), AF488- or APC-eFluor780-conjugated anti-CD24 (M1/69), BB700-conjugated anti-NK1.1 (PK136), PECy7 or AF488-conjugated anti-CD3∊ (145-2C11), APC- or AF700-conjugated anti-TCRβ, PE-Cy7-conjugated anti-CD44 (IM7), PE-, BV421- or PECy7-conjugated anti-CD138 (281-2), APC-, SB600- or PerCp-Cy5.5-conjugated anti-Sca-1 (D7), BV786- orAPC-Fire750-conjugated anti-FR4 (GK1.5), SB702-conjugated anti-CD19 (1D3) and APC-Fire750-conjugated anti-CD4 (GK1.5). Dead cells were stained with LIVE/DEAD® Fixable Aqua Dead Cell Stain kit (ThermoFisher Scientific, Illkirch, France). PBS-57 glycolipid-loaded and -unloaded control CD1d tetramers (BV421 or PE-conjugated) were from the NIH Tetramer Core Facility (Emory University, Atlanta, GA). For intracellular staining, Pacific Blue or APC-conjugated anti-IFN-γ (XMG1.2), PE-conjugated anti-IL-4 (11B11), BV650-conjugated anti-IL-17A (TC11-18H10), FITC-conjugated anti-mKi67 (11F6), AF488- or APC-conjugated anti-PLZF (R17-809), PE-CF594- or APC-conjugated anti-RORγt (Q31-378), PE-Cy7- or PE-conjugated anti-T-bet (4B10), MitoprobeTM TMRM and APC-conjugated Mitotracker (ThermoFisher Scientific, Illkirch, France) were used. Phorbol 12-myristate 13-acetate (PMA) and ionomycin were from Sigma-Aldrich (Saint-Quentin-Fallavier, France). Mouse ELISA kits were purchased from R&D systems (Lille, France). α-Galactoslcermaide (α-GalCer) was from Coger (Paris, France).

### Tissue preparation and flow cytometry

Single-cell suspensions were prepared from thymus, liver and spleen as previously described(Paget et al., 2007). Red blood cells were removed using a red blood cell lysis buffer (Sigma-Aldrich). In some cases, cells were stimulated for 4 hours in complete media containing PMA (100 ng/ml) and ionomycin (1 μg/ml) in presence of protein transport inhibitor cocktail (eBioscience) added 30 min after stimulation. For intracellular staining, cells were first stained with antibodies to surface makers and viability dye (LIVE/DEAD Fixable Aqua Dead Cell Stain), then fixed and permeabilized using a Fixation/Permeabilization Solution Kit (BD Biosciences) or a True-Nuclear Transcription Factor Buffer Set (Biolegend) for intracytoplasmic intranuclear staining respectively. Events were acquired on either LSR Fortessa (BD Biosciences) or a MACSQuant (Miltenyi Biotec) cytometers. Analyses were performed by using the VenturiOne software (Applied Cytometry; Sheffield, UK) or FlowJo v10 (BD Biosciences).

### Single-cell RNA-seq and data pre-processing

Single cell suspension from 4 thymi were pooled and total iNKT thymocytes (CD45^+^ CD3^+^ PBS57-loaded CD1d tetramer^+^) were sorted into non-HEPES medium containing 10% of FCS. Sorted cells were counted under a microscope and 10,000 cells were loaded onto a Chromium 3’ chip. Reverse transcription and library preparation were performed according to the manufacturer’s protocol (10X Genomics). Libraries were sequenced on a NextSeq 500 High Output v2 kit (75 cycles) that allows up to 91 cycles of paired-end sequencing: read 1 had a length of 26 bases that included the cell barcode and the UMI; read 2 had a length of 57 bases that contained the cDNA insert; Index reads for sample index of 8 bases. Cell Ranger Single-Cell Software Suite v2.0.0 was used to perform sample demultiplexing, barcode processing and single-cell 3′ gene counting using standards default parameters and *Mus musculus* build mm10.

### Single-cell RNA-seq analysis

#### Cell and gene filtering

Raw gene expression matrices generated per sample were loaded and processed within Scanpy python workflow(Wolf et al., 2018). Cells having more than 5% of mitochondrial RNA were removed; then ribosomal, mitochondrial and genes expressed in less than 3 cells were removed.

#### Doublet removal

Unbiased identification of doublets (technical error) in samples was performed using Doublet Detection and Scrublet (https://github.com/JonathanShor/DoubletDetection, https://github.com/AllonKleinLab/scrublet). Using default parameters, we removed the cells predicted as doublets by any of the two methods.

#### Statistical analysis and visualization

Raw counts of remaining cells were normalized and log-transformed before highly variable genes (HVG) was identified using Scanpy HVG function (Parameters: min_mean=0.0125, max_mean=3, min_disp=0.5). Resulting HVG were summarized by principle component analysis, then the first 50 principle components were further summarized using UMAP dimensionality reduction. Cells were then clustered using the Louvain algorithm with a resolution of 1.5. Cell clusters in the resulting UMAP two-dimensional representation were annotated to known biological cell types using canonical marker genes identified using Scanpy rank_gene_group function with the Wilcoxon’s rank test.

#### Cell cycle Analysis

Cell cycle scoring (S phase and G2M phase) was performed using Scanpy score_genes_cell_cycle function with cell cycle genes list from Tirosh et al. (https://science.sciencemag.org/content/352/6282/189).

#### Data integration

For *Fhl*2^+/+^ and *Fhl2*^−/−^ single cell transcriptome comparison, batch effect was removed using mutual nearest neighbours method implemented in ‘mnn_correct’ function in mnnpy python package (Haghverdi L) using intersection of individual sample highly variables gene sets.

#### Trajectory inference using PAGA

For each of the two samples, cycling iNKT and NKT1d clusters was removed before independent trajectory inference analysis was performed using PAGA algorithm available within Scanpy.

#### Functional enrichment

Data gene sets were analyzed for functional enrichment with Enrichr (https://amp.pharm.mssm.edu/Enrichr/#) using as the reference databases GO biological process 2018, GO molecular function 2018 and KEGG 2019 mouse.

### Data and Code availability

All data are currently submitted to the Gene Expression Omnibus accession number GSE141895. Data is also available through a dedicated web interface (https://www.genomique.eu/cellbrowser/ucagenomix/baranek2020/). All Pyhton and R scripts used for single cell statistical analysis can be accessed on github (https://github.com/ucagenomix/sc.baranek.2020).

### Artificial thymic retention

Mice received daily 20 μg of FTY720 (Sigma-Aldrich) intraperitoneally (i.p.) for seven days. Then thymi of treated mice were analyzed the day following the last administration.

### *In vivo* iNKT cell activation

Mice were injected i.p. with 2 μg of α-GalCer (or vehicle) and spleen and liver were harvested 8 hours post-injection.

### Reaggregate Thymus Organ Cultures (RTOC)

B6 Thymocytes were stained with PE-conjugated PBS57-loaded CD1d tetramers and enriched using Mojo Sort anti-PE nanobead (Biolegend). Then, iNKT subsets were sorted using a BD FACS Melody (BD biosciences). 2 x 10^4^ purified iNKT cells were cocultured with 2 x 10^7^ of total thymocytes from *Traj18*^−/−^ mice in 24-well culture plate in complete 10 % FCS RPMI1640 containing recombinant IL-7 (20 ng/mL) and recombinant IL-15 (20 ng/mL) (Miltenyi Biotech). Cells were incubated for 4 days and dead cells were removed by Ficoll-Paque PLUS (Fisher scientific) centrifugation prior analysis.

### Statistical analysis

All statistical analysis was performed by using GraphPad Prism software. The statistical significance was evaluated by using nonparametric unpaired Mann-Whitney *U* tests in order to compare the means of biological replicates in each experimental group. Results with a P value of <0.05 were considered significant. ns: not significant; *p < 0.05; **p < 0.01; ***p < 0.001.).

---

**Supplemental Table 1:** Top-200 differentially up-regulated genes per cluster in WT iNKT cells.

**Supplemental Table 2:** List of iNKT0 marker genes encoding for products involved in nucleic acid regulatory processes and epigenetic landscape.

**Supplemental Table 3:** Top-200 differentially up-regulated genes in iNKT2a vs iNKT2b clusters.

**Supplemental Table 4:** List of marker genes in subclusters of cycling iNKT (**a**) and iNKT1d (**b**).

**Supplemental Table 5:** List of differentially regulated genes in *Fhl2*^+/+^ vs *Fhl*2^−/−^ iNKT cells per cluster. **a**, total iNKT cells; **b**, iNKT2a; **c**, iNKT2b; **d**, iNKT1b; **e**, iNKT1c; **f**, iNKT17.

