## Supplemental Figures for "High dimensional single-cell analysis reveals iNKT cell developmental trajectories and effector fate decision"

**Baranek et al.,**

**Supplementary data**

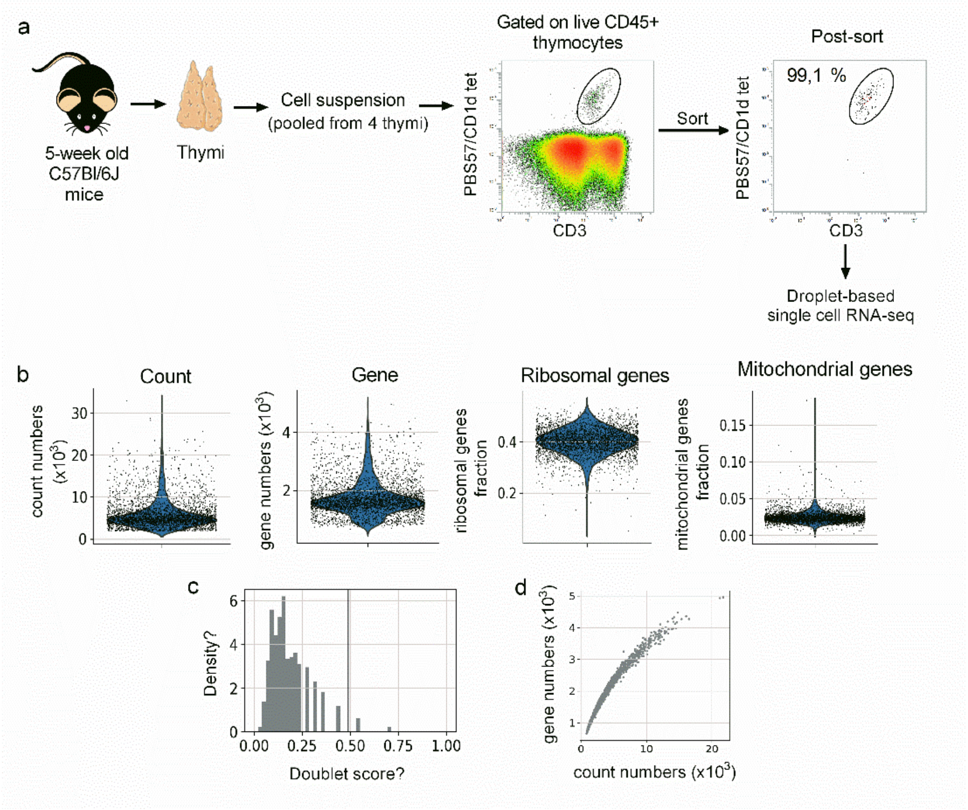


**Figure S1: scRNA-seq of total thymic iNKT. a,** Schematic representation of the experimental workflow used to generate scRNA-seq data from total iNKT thymocytes. **b**, Scatter plots showing quality control metrics (counts, genes, ribosomal and mitochondrial genes). The violin plot represents the density of the continuous variable for each value. Each dot is representative of one cell. **c,** Density histogram showing the numbers of UMIs detected per cell. The grey line represents the threshold used to eliminate doublet. **d,** Scatter plot showing the number of cells per gene.

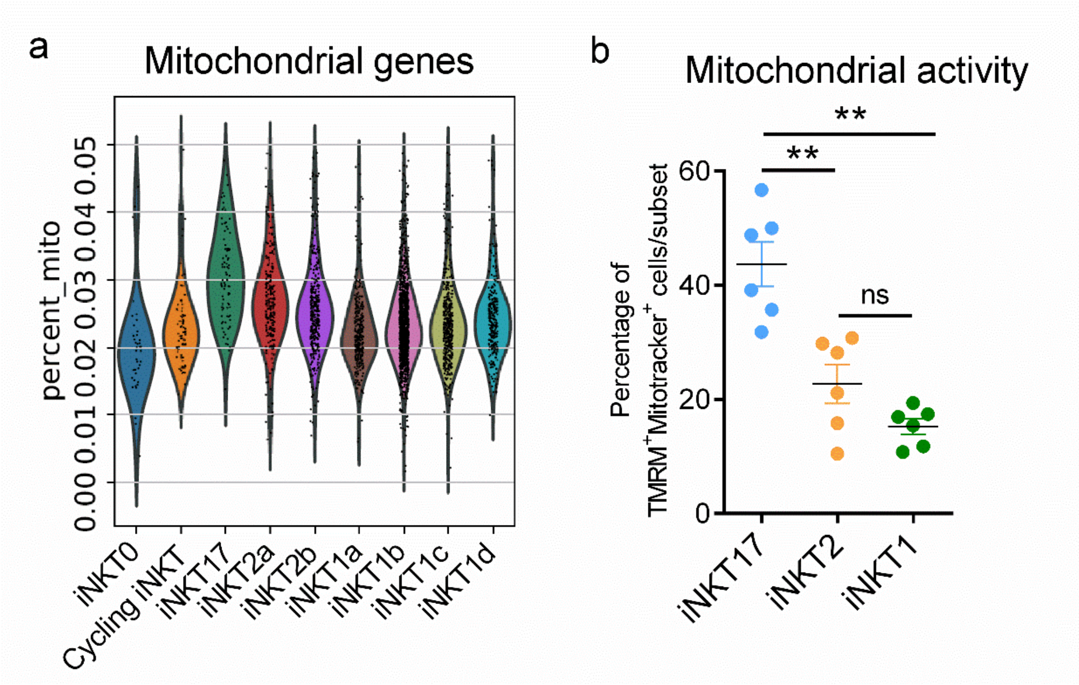


**Figure S2: Mitochondrial activity in iNKT thymocyte subsets**. **a**, Percentage of mitochondrial RNAs detected in each iNKT subsets as identified in Fig. 1a. **b,** Flow cytometric analyses of mitochondrial activity of iNKT thymocytes in the thymus of 5 week-old C57BL/6J mice. Individual values and means ± SEM of $TMRM^{+}Mitotracker^{+}$ cells for each subsets are depicted in the right panel. Data are representative of one experiment out of three. **, p<0.01

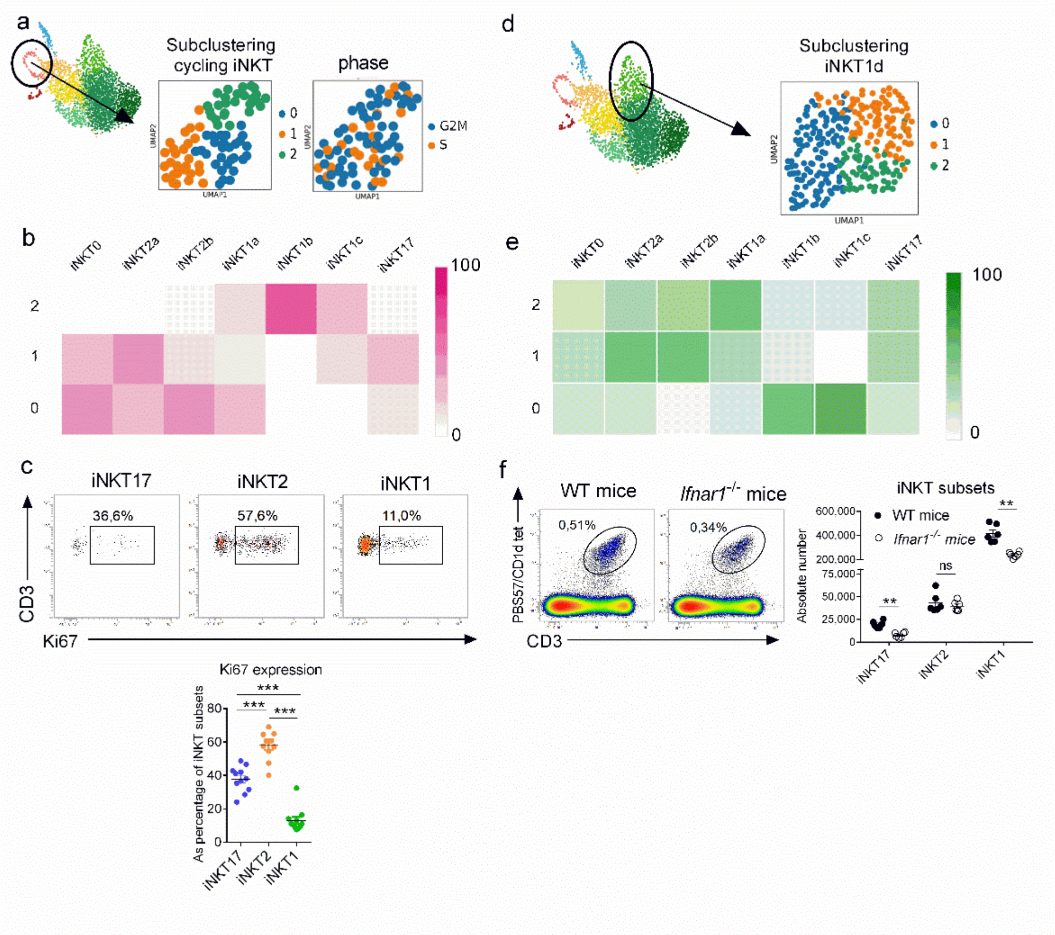


**Figure S3: Particular biological activity guided the clustering of cycling iNKT and iNKT1d. a,** Clustering of cycling iNKT using the top 50 DEGs in the other seven iNKT clusters. UMAP visualization of the clusters (left) and of the quantitative scores for G2M and S phases for each cell (right) obtained. **b,** Heatmap comparing the three clusters obtained in a. with each iNKT subsets using the Morpheus software (https://software.broadinstitute.org/morpheus). **c,** Flow cytometry showing Ki67 expression in iNKT1, iNKT2 and iNKT17 subsets. Data are representative of three independent experiments. Individual values and means ± SEM of mean fluorescence intensity (MFI) for each subsets are depicted in the lower panel. **d,** Clustering of iNKT1d using the top 50 DEGs in the other seven iNKT clusters. UMAP visualization of the clusters obtained. **e,** Heatmap comparing the three clusters obtained in d. with each iNKT subsets using the Morpheus software. **f,** Representative dot plots showing total iNKT cells (live $CD3^+$ PBS57/CD1d $tetramer^+$ cells) from 5 week-old *Ifnar1*$^{-/-}$. Individual values and means ± SEM of absolute numbers of iNKT1, iNKT2 and iNKT17 subsets are presented in the right panel. Data are from one representative of two experiments. **, p<0.01; ***, p<0.001

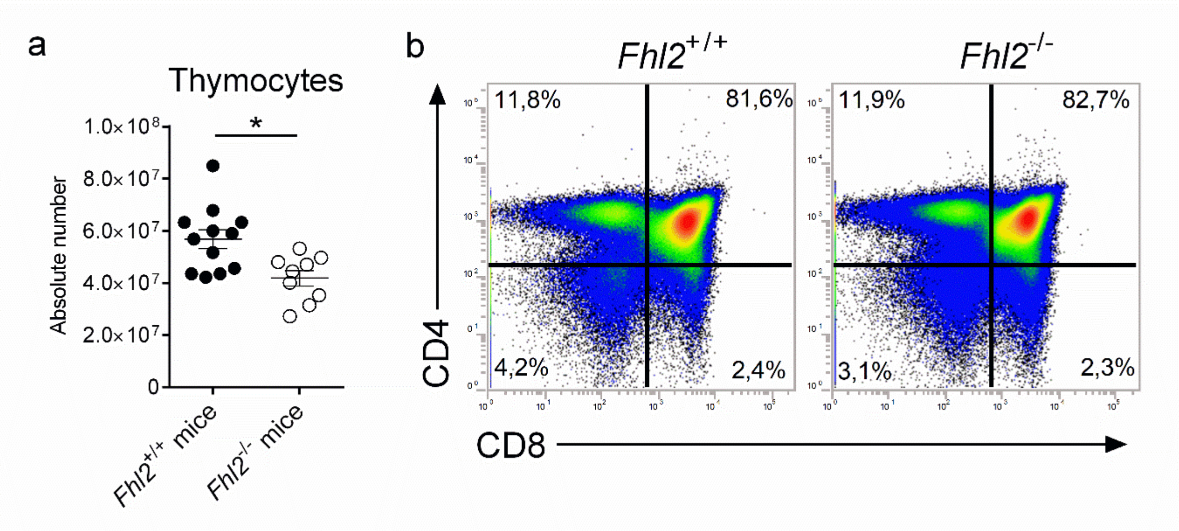


**Figure S4: Thymocyte subsets in *Fhl2*$^{-/-}$ mice**. **a**, Individual values and means ± SEM of absolute numbers of thymocytes are depicted. Data are from two experiments. *, $p<0.05$. **b,** Flow cytometry showing CD4 and CD8 expression on thymocytes in *Fhl2*$^{+/+}$ and *Fhl2*$^{-/-}$ littermate mice. Representative dot plots from two experiments are shown.

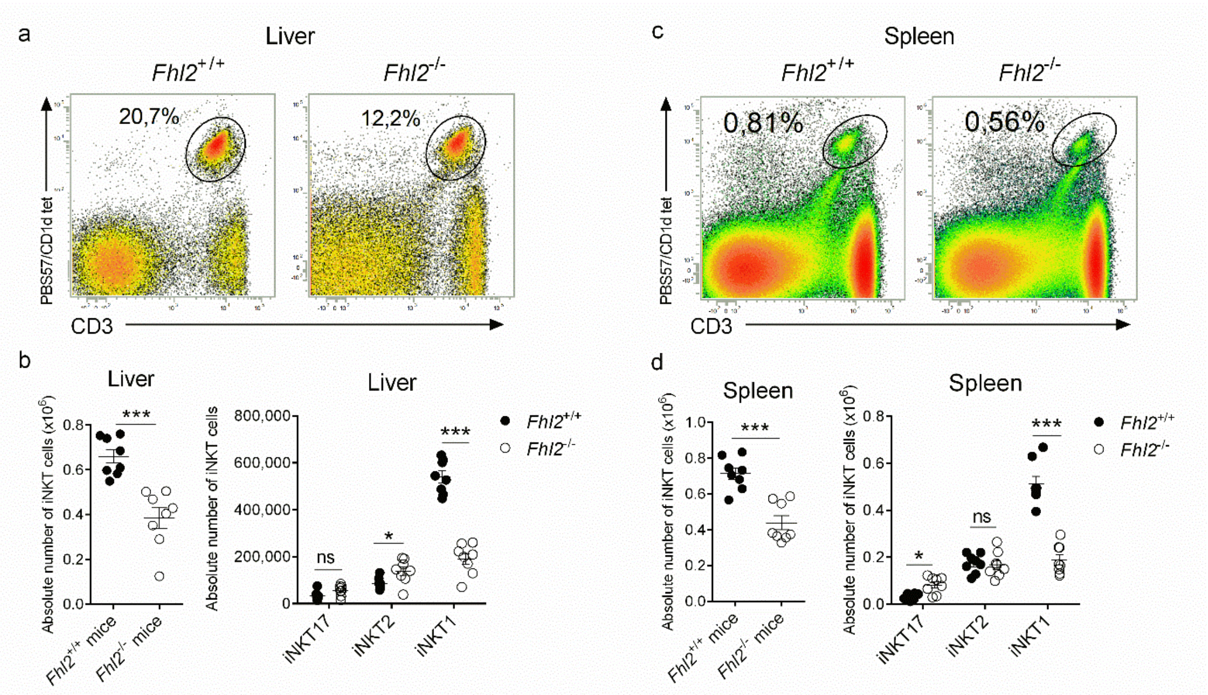


**Figure S5: iNKT cell subsets in peripheral tissues of *Fhl2*$^{-/-}$ mice**. Flow cytometric analyses of iNKT subsets in the liver (a-b) and the spleen (c-d) collected from 5 week-old *Fhl2*$^{+/+}$ and *Fhl2*$^{-/-}$ littermate mice. **a, c**, Representative dot plots showing total iNKT cell (live CD45$^{+}$ CD3$^{+}$ PBS57/CD1d tetramer$^{+}$ cells) percentages. **b, d**, Individual values and means ± SEM of absolute number of total iNKT cells (left) and of iNKT1, iNKT2 and iNKT17 subsets (right) are presented. ***, p<0.001.
